# Traditional Norwegian Kveik Yeast: An Ancient Sister Group to Domesticated *Saccharomyces cerevisiae*

**DOI:** 10.1101/2023.07.03.547515

**Authors:** Michael Dondrup, Hans Geir Eiken, Atle Ove Martinussen, Lisa Karine Haugland, Rita Holdhus, David Dolan, Sushma Grellscheid, Snorre Hagen, Abdelhameed Elameen, Tor Myking

## Abstract

Kveik is the common name of yeast that has been used in traditional farmhouse brewing of western Norway for generations. Its fast fermentation, increased flocculation, temperature tolerance, and rich flavor profile have led to growing interest in recent years. Previous genetic analyses have shown that kveik forms a distinct group within the *Saccharomyces cerevisiae* tree and placed its origins within the Beer I clade of industrial brewing yeasts, although with signs of mixed ancestry.

In this study, we revisited the phylogenetic position of kveik within the *S. cerevisiae* tree. We searched for traditional farm breweries in western Norway and collected ten samples of potential kveik yeast.

Using Illumina whole genome shotgun sequencing, we reconstructed the phylogenetic tree of kveik based on *de novo* genome assemblies and variant calls of our new kveik samples, along with published wild and domesticated *S. cerevisiae* strains. We calibrated and used sequential computational experiments at different thresholds to determine the most probable phylogenetic position of kveik yeast.

Previously sequenced kveik genotypes form a clade with our new samples clustering partially by place of origin. Our results indicate that kveik is indeed a compact clade within *S. cerevisiae* with significantly reduced polymorphism compared to common brewing yeasts and wild strains. Contrary to what was previously thought, our analyses support a more ancient divergence of kveik and place it closer to the root of the *S. cerevisiae* tree.

In conclusion, our genetic analyses suggest that kveik is a unique and ancient yeast group, distinct from other domesticated *S. cerevisiae* strains. Considering a possible far east origin of kveik yeast, the apparent endemism to western Norway remains as a big paradox These findings have important implications for the understanding of yeast domestication and the use of kveik in modern brewing practices.

## Introduction

The budding yeast *Saccharomyces cerevisiae* has been used by humans for thousands of years to ferment various food and beverage products. Recent phylogenetic data indicate that East Asia and China may be the center of origin of one or several domestication events (Duan et al. 2018; Peter et al. 2018). Genome analyses have revealed that wild and domestic populations of *S. cerevisiae* are separate groups, and furthermore that there are two distinct domesticated groups linked to liquid and solid-state fermentations that both originate from a single ancestor (Duan et al. 2018; Han et al. 2004; Peter et al. 2018).

Within *S. cerevisiae* strains, hallmarks of domestication appear most pronounced in beer yeasts with reduced survival in nature, reduced sexual reproduction and stress resistance, as well as increased maltotriose utilization and genome decay (Gallone et al. 2016). The reason for this is evolution in a highly selective niche where beer yeasts are harvested and re-used after the fermentation process to start a new fermentation (Gallone et al. 2018). Phylogenetic results divide industrial beer yeast into two major clades where the Beer 1 clade shows the strongest signs of domestication and geographical boundaries, while the Beer 2 clade shows more limited domestication (Gallone et al. 2016). Modern industrial brewing yeasts may have their last common ancestor around 1600-1700AD before the era of modern microbiology (Gallone et al. 2018; Gallone et al. 2016), but a long time after the estimated invention of beer brewing at 3000 BC (Gallone et al. 2016; McGovern et al. 2004; Michel et al. 1992). In line with this, analysis of a sample of human paleofeces from the Hallstatt mine works from the iron age (dated 650-545 BC) identified abundant *S. cerevisiae* DNA. The sample contained ancient DNA of a strain related to the Beer 2 clade that was likely used for alcoholic fermentation (Maixner et al. 2021). Still, it seems that important parts of the evolutionary history and diversity of beer yeasts remain to be unraveled. Today, there are over 2500 sequences genomes of wild and domesticated strains of *S. cerevisiae* (Libkind et al. 2020). This wealth of data has the potential to aid in the development of new fermentation processes and applications for *S. cerevisiae,* as well as shed light on its evolutionary history and domestication (Libkind et al. 2020).

Farmhouse brewing has a longstanding tradition in Norway and remains a vibrant part of the local culture to this day at the west-coast of Norway (Garshol 2023). The yeast used in farmhouse brewing was usually cultivated and passed down through generations within the same family. On the west coast of Norway this type of yeast is named “kveik”. There is growing interest in kveik yeasts by brewers worldwide due to its reported properties such as fast fermentation, strong flocculation, high temperature and ethanol tolerance, and the production of multiple flavor metabolites with little off-flavors at high temperatures (Kawa-Rygielska et al. 2022; Kits and Garshol 2021; Preiss et al. 2018; Foster et al. 2022). There are differences between the different kveik stains regarding fermentation temperature range, but all show an increased fermentation rate compared to beer yeasts (Foster et al. 2022). The disaccharide trehalose is known to protect yeast from temperature and ethanol stress, and kveik yeasts accumulate trehalose faster than beer yeasts, which may be caused by a mutation in the Trehalose Synthase Complex (Foster et al. 2022).

Recently, genetic analysis indicated that six kveik strains formed a distinct group of domesticated *S. cerevisiae* yeast within the Beer 1 clade (Preiss et al. 2018). However, the findings were not clear as there were signs of a mixed ancestry with possible unknown parent strains. Also, one “kveik-culture” was found to be an interspecies hybrid with *Saccharomyces uvarum* (Krogerus, Preiss, and Gibson 2018). Thus, there is a need for more knowledge on the genetic composition, origin, and phylogeny of kveik yeast.

The historical tradition, the phenotypes and genotypes of the different stains all indicate that this rare type of *S. cerevisiae* yeast may be essential to understand the domestication process of beer yeast in general. The partially unclear ancestry of kveik led us to use genome analysis to revisit the phylogenetic relationship of kveik yeast to other domesticated and wild *S. cerevisiae.* Our first aim was to collect kveik yeast in its original form from items used in the old farm brewing traditions in western Norway. Next, we aimed at developing harmonized whole genome alignments of kveik to domesticated and wild groups of yeasts to investigate both the possible age and the phylogenetic position of kveik yeasts relative to beer and wine yeasts. To achieve this goal, we intended to validate our bioinformatic analysis through multiple computational methods and experimental tests. Our overall aim was to gain knowledge on the possible geographic and phylogenetic origins of kveik.

## Materials and methods

### Sampling

Traditional farmhouse brewing has long traditions in western Norway and is still actively practiced. Through knowledge of these traditions and relevant networks in the region we were able to locate brewers and document brewing traditions from previous generations. Important criteria for sampling were: 1. Kveik yeast had been in the family for several generations. 2. The intangible cultural heritage connected to brewing with kveik yeast was still practiced. 3. The kveik yeast had recently been used for brewing. The tradition bearers were willing to be interviewed regarding the processes of brewing and the use of kveik yeast. Ten samples were collected directly from owners brewing equipment with a written contract of use in research (Table 1 and Figure 1, red dots). Sample 1 was re-sampled after two years from the same owner (sample 1A and 1B). In addition, six genomes from kveik strains from western Norway previously isolated by Preiss et al. 2018 were used for alignments (blue dots in Figure 1) and of these, two samples were from the same location and owner (Table 1).

**Figure 1:**
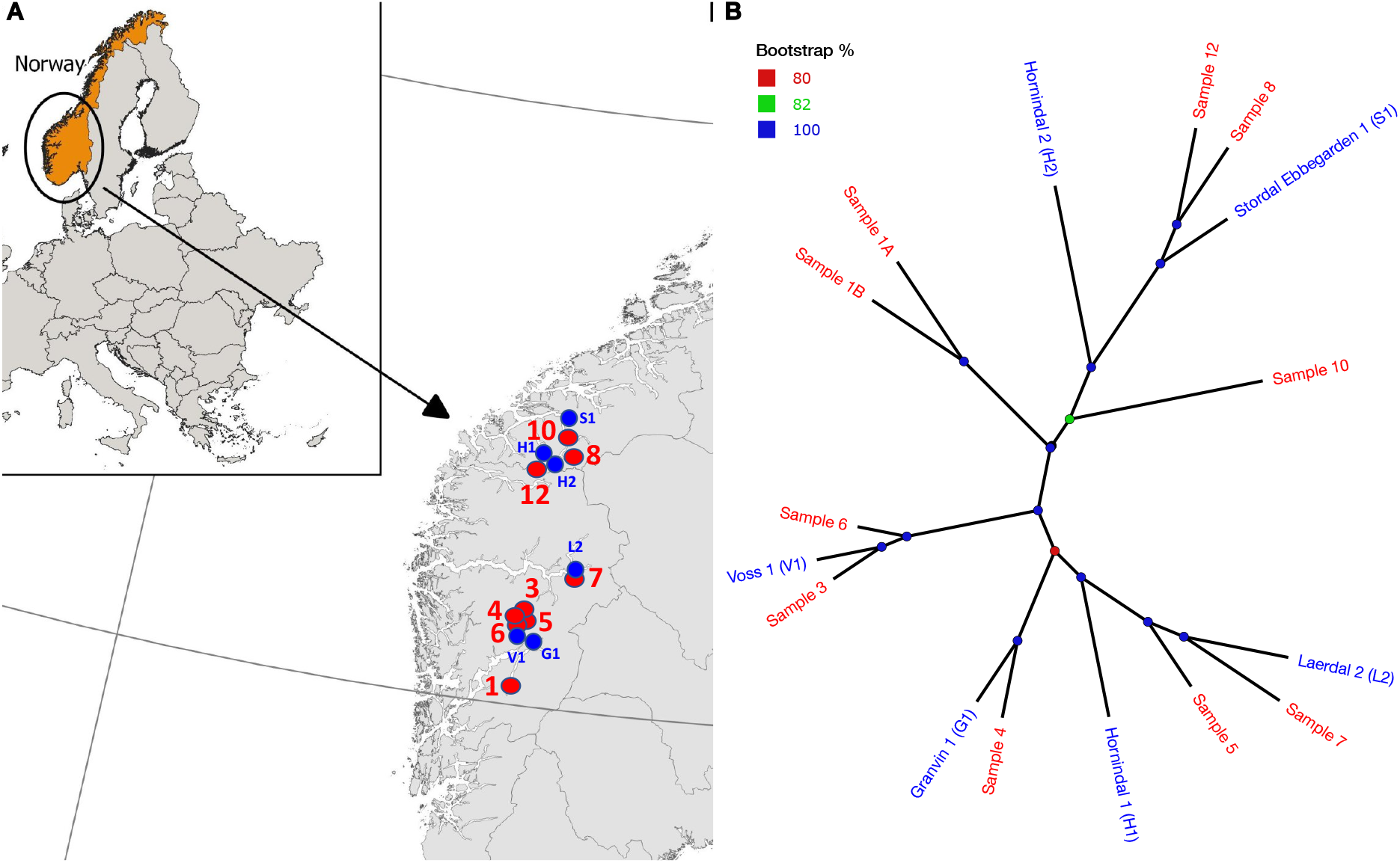
A) Geographic origin in western Norway of kveik yeast strains analyzed in this study (red labels and dots). Blue dots show the geographical location of six strains previously isolated (Preiss et al. 2018). Sample number 6 and 7 were collected from the same location and owner as in the previous study (indicated by partial overlapping red and blue dots). B) Unrooted ML phylogenetic tree of the kveik clade (from Tree 1, Fig. 2A), label colors as in A); node support indicated as bootstrap % by node color.

**Figure 2:**
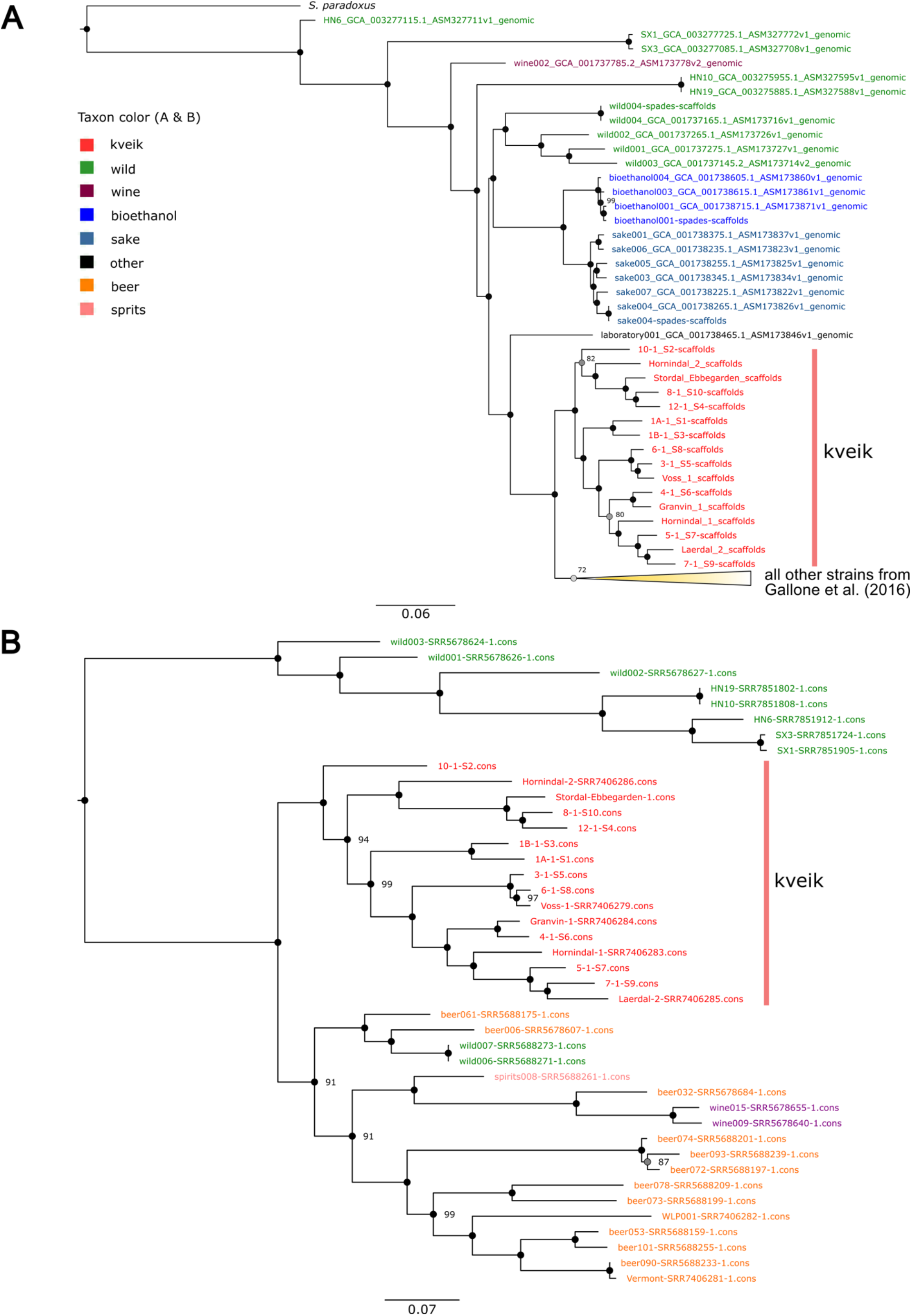
Phylogenetic analysis of Kveik. A) ML phylogenetic tree from de novo assemblies B) ML tree based on filtered variant calls from 41 strains using the REF allele. Scalebars in substitutions per site. Kveik groups in A, C, and D are marked in red. Tree A) is rooted on the S. paradoxus genome, B) is rooted on the outgroup of wild taxa. Node bootstrap support is given in percent only for nodes with support < 100%. Taxa having suffix “scaffolds”: result from assembly with SPAdes, “genomic”: downloaded from GenBank, “cons”: result from BCFtools consensus method using the REF allele in heterozygous loci.

**Table 1:** Sample origin of traditional kveik yeast. Contact details of the owners can be made available upon reasonable request.

| Sample ID | Location | Geo-location (EUREF 89 UTM 33) | Form | Last used in brewing | Collected from same owner in previous study by Preiss et al. 2018 (GeneBank No.) |
| --- | --- | --- | --- | --- | --- |
| 1A | Odda | N6692324<br>E29532 | Liquid | 2020 |  |
| 1B | Odda | N6692324<br>E29532 | Dry | 2018 |  |
| 3 | Vossestrand | N6768607<br>E44635 | Dry | 2020 |  |
| 4 | Dyrvedalen, Voss | N6754153<br>E25933 | Liquid | 2020 |  |
| 5 | Bordalen, Voss | N6747804<br>E33831 | Liquid | 2020 |  |
| 6 | Voss | N6753313<br>E24365 | Liquid | 2020 | MG641169 |
| 7 | Lærdal | N6790637<br>E101732 | Dry | 2021 | MG641176 |
| 8 | Geiranger | N6906725<br>E95639 | Dry | 2021 |  |
| 10 | Eidsdal | N6921101<br>E92777 | Dry | 2021 |  |
| 12 | Grodås | N6899623<br>E56769 | Dry | 2021 |  |

### Yeast cultivation and single colony isolation

Dried kveik samples were rehydrated in sterile water. The rehydrated samples of kveik and the liquid yeast slurries were enriched by inoculating 100 μl of the slurry into 5 ml YPD (1% yeast extract; 2% peptone; 2% dextrose; chloramphenicol 0.3 g/l). The samples were incubated at 30℃ for 24 h with shaking, then streak plated onto Wallerstein Nutrient agar (with chloramphenicol 0.3g /l) (WLN; Thermo Fisher CM0309), a differential medium for yeasts that distinguishes multiple yeasts from each other within one sample based on uptake of the bromocresol green dye. Yeast colonies were then sub streaked two times onto WLN to ensure purity. The single colonies were enriched for DNA extraction as described above. The remaining overnight cultures were long-term stored in aliquots at minus 80℃ using a glycerol-based standard storage medium (peptone 1% w v-1, yeast extract 0.5% w v-1, glucose 1% w v-1, glycerol 25% v v-1).

### DNA extraction

DNA extraction was performed based on a modified protocol following (Preiss et al. 2018) in combination with Qiagen DNeasy Plant Mini kit. In short, yeast cells were collected from 1.5 ml of the overnight yeast culture by pelleting cells with centrifugation at 13000 rpm for 5 min, discharged the supernatant, washed the cell pellet in 1 mL of 0.8% NaCl and added 0.3 g of 0.5-mm-diameter glass beads. The cell-bead mixture was centrifuged at 11600 x g for 10 min at 4°C, and the supernatant were discharged followed by resuspension in 300 μl of breaking buffer (2% Triton X-100, 1 % SDS, 100mM NaCl, 10 mM Tris-HCl) and 300 μl of phenol-chloroform-isoamyl alcohol (25:24:1). The suspension was vortexed at high speed 3 x 45 sec, centrifuged at 11600 x g for 10 min at 4°C, and the final aqueous phase was then used for DNA extraction with the DNeasy Plant Mini Kit (Qiagen, Tokyo) according to the manufacturer’s instructions.

### Genome sequencing

DNA samples were quantified using Qubit and 100ng of each sample was used for library construction applying Illumina TruSeq DNA nano prep. Quantification of libraries was done by using KAPA Library Quantification kit (QPCR) for Illumina, and average library input fragment size was estimated using on Agilent TapeStation. Sequencing was performed on the Illumina NovaSeq 6000 Sequencing System using standard workflow and sequencing parameters at 2×150 bp pair-end sequencing with an average coverage of 10000x per kveik yeast genome.

### Variant analysis

Raw data from Preiss et al. (2018) and Gallone et al. (2016) were downloaded from the Sequence Read Archive (SRA). Raw data in FASTQ format were quality controlled with FastQC and MultiQC (Ewels et al. 2016) before and after trimming. All reads were trimmed with Trimmomatic (Bolger et al. 2014). Reads from (Preiss et al. 2018) contained some Illumina universal adapter sequences that were not properly removed by Trimmomatic and all samples were therefore subjected to a second round of adapter trimming with cutadapt (Martin 2011). Reference-based multiple sequence alignment and variant calling were performed using the *S. cerevisiae* RefSeq genome R64-2-1 (GCF_000146045.2) (Engel et al. 2022). Gene descriptions were downloaded from Ensembl BioMart (Cunningham et al. 2021).

*De novo* variant detection was then carried out by aligning the trimmed reads against the reference genome with SpeedSeq (Chiang et al. 2015) and the resulting BAM files were filtered by SAMtools (Danecek et al. 2021) retaining alignments with minimum mapping quality 50. Variant analysis was done with The Genome Analysis Toolkit (GATK) v4.2.6.1 (Mckenna et al. 2010) using HaplotypeCaller with ploidy parameter set to 4 and 2 and FreeBayes v1.1.0 (Garrison and Marth 2012) with ploidy = 4. Duplicate alignments were first marked using GATK MarkDuplicatesSpark for both tools.

Joint genotypes were called by GATK GenomicsDBImprot → GenotypeGVCFs exluding the mitochondrial genome (Depristo et al. 2011). The resulting variants were then filtered using GATK VariantFiltration with the following filters: mapping quality (MQ) < 40, minimum depth (DP) < 5, quality by depth ratio (QUAL / DP) < 2.0, genotype quality (GQ) < 30, setting filtered genotypes to no call and retained only biallelic SNPs and InDels that passed all filters. The output of GATK HaplotypeCaller with ploidy setting 2 was used only for association testing and calculating windowed statistics. Genome-wide nucleotide diversity and neutrality (Pi and Tajima’s D) was estimated with VCFtools (Danecek et al. 2011) with window size =10,000 bases and then calculating the arithmetic mean over all bins. Association testing was performed in PLINK v1.9 (Purcell et al. 2007). Therefore, unfiltered variants (Ploidy = 2) were annotated with SnpEff (Cingolani et al. 2012) and then filtered to retain only missense variants. The assignment to either “kveik” or “beer” group was used as phenotype. Bonferroni correction was applied by multiplying p-values with the number of tests performed.

Variants were annotated for potential effect with SnpEff (Cingolani et al. 2012). Gene Ontology (GO) enrichment analysis of genes highly affected by variants were based on the predictions of variant impact and carried out in the Saccharomyces Genome Database (SGD) (Engel et al. 2022). Copy Number Variation (CNV) analysis was carried out using control-FREEC (Boeva et al. 2011) without specifying a reference and ploidy = 4. Variant files were visualized and visually inspected with the Integrative Genomics Viewer (IGV) (Robinson et al. 2011).

### Variant analysis of selected genes

Consensus sequences for single genes were generated from the aligned BAM files using SAMtools (Danecek et al. 2021). Sequences were further clipped, translated, analyzed, and visualized in Jalview (Troshin et al. 2011; Waterhouse et al. 2009). Multiple sequence alignments of the consensus nucleotide sequences and the reference were generated with Muscle (Edgar 2004) with default settings. Multiple sequence alignments of amino-acid sequences including stop-codons were generated in ClustalOmega (Sievers et al. 2011). Protein tertiary structure of RPI1 as predicted by AlphaFold (Varadi et al. 2021) was downloaded from UniProtKB (The UniProt Consortium 2022) and visualized in Chimera (Pettersen et al. 2004).

### Genome assembly and phylogenetic analysis

Trimmed reads were *de novo* assembled using the short read assembler SPAdes (Bankevich et al. 2012). Reference-based alignment of genome assemblies was performed using the NASP pipeline (Sahl et al. 2016) with all default parameters. NASP uses NUCmer to align assembled genomes (Delcher et al. 2003). To generate a phylogeny, we generated a filtered SNP matrix retaining only sites overlapping a coding sequence (as annotated by SnpEff) and retaining only parsimony-informative sites (minor allele count > 1) leading to Tree 1 and also the matrix of genome-wide SNPs (bestsnps.fasta) generated by NASP leading to Tree 2. IQ-Tree2 v2.1.2 (Nguyen et al. 2015; Minh et al. 2020) was used with 1000 iterations of the ultrafast bootstrap approximation (Minh et al. 2013) using ModelFinder for model selection (Kalyaanamoorthy et al. 2017); best-fit models chosen by ModelFinder according to BIC: TVMe+ASC+R4 (Tree 1, CDS only) and TVM+F+ASC+R4 (Tree 2, bestsnps.fasta). The resulting trees were visualized with FigTree (Rambaut 2023) and the *Saccharomyces paradoxus* genome (Yue et al. 2017) was used as outgroup to root the trees. The branch length of *S. paradoxus* was scaled by a factor of 0.01 in Tree 2 for improving visualization with the genome-wide SNP matrix. Principle component analysis (PCA) of the alignment in the SNP-matrix was computed in Jalview (Waterhouse et al. 2009). Projections of input vectors on the first two principal components were plotted in R (R Core Development Team 2022).

### Phylogenetic analysis based on variants

A series of experiments was carried out using raw sequencing data of 41 strains downloaded from SRA. Filter and selected variants from GATK (ploidy = 4) were then further filtered at different MAF thresholds (0.01, 0.05, 0.1, 0.2, and 0.3) and alternative consensus genotypes were generated with BCFtools (Danecek et al. 2021) to retrieve consensus genomes with either only ALT, or REF alleles (leading to Fig. 3B) inserted at polymorphic sites and in one case we retained only variants overlapping genes. The alternative genotypes were then aligned in NASP, and phylogenetic trees and consensus topologies were generated with IQ-Tree2 as described above.

**Figure 3:**
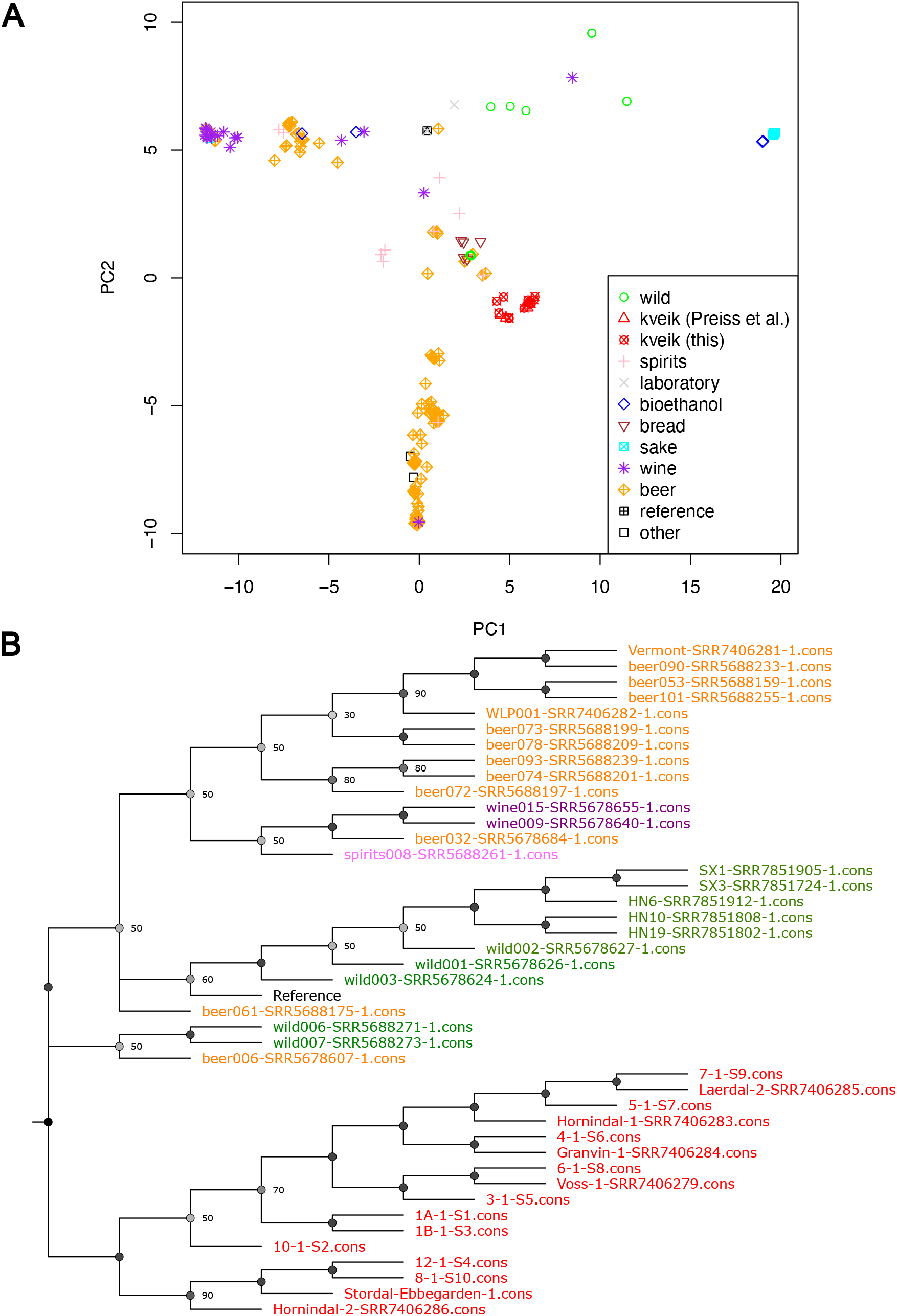
A) Principal component analysis of the variant matrix (bestsnp.fasta) with points projected on the first two principal components (PC), S. paradoxus at (-10.775667, -56.893497) not shown. Percent variance explained: PC1: 95.17%, PC2: 0.6% B) Mid-point rooted consensus topology of 10 trees based on filtered GATK variants using either REF and ALT alleles and different MAF cutoffs (0.01, 0.05, 0.1, 0.2, 0.3). Node consensus support is given in percent only for nodes with support < 100%. For nodes with support ≤ 50%, no absolute majority exists.

### Calibration experiments and reproducibility

To evaluate whether we could use the workflow using compound consensus sequences, we first conducted a series of experiments to assess robustness and reproducibility. We created a workflow very similar to Preiss et al. (2018) except for program versions. Variants from the freebayes analysis (ploidy = 4) were filtered for variants with MAF < 0.01. We generated alternative genotypes using the ALT allele with BCFtools. The alternative genotypes were then analyzed jointly in the NASP pipeline with the genome assemblies by (Gallone et al. 2016) including those assemblies for which we also had generated variant-based genotypes. The resulting bestsnp.vcf file was annotated with SnpEff and filtered as follows: only bi-allelic SNPs that were informative (different genotype in at least 2 samples, equivalent to filtering “MAF > 0.01” by Preiss et al.) and overlap with a protein-coding gene region, using custom Perl/shell scripts. The resulting SNP matrix was then used with IQ-Tree2 using the nucleotide substitution model GTR+R4. The same analysis was run with three different datasets: 1) Containing the same data as in Preiss et al. (2018), 2) adding all our samples except sample 8 and 5 selected samples from Gallone et al. (2016), and 3) all 41 strains from the variant analysis.

## Results

We collected 10 samples of potential kveik yeast from owners in western Norway (Table 1, Figure 1, red dots), isolated single yeast colonies and subjected the samples to whole genome sequencing. Deep genomic sequencing of approximately 1000X (Table 2) was followed by *de novo* genome assembly.

**Table 2:** Per-sample statistics of filtered GATK variant calls and de novo assembly. Variants on the mitochondrial genome were excluded.

| Sample ID | # Read pairs | Contig N50 | Scaffold N50 | # NonRef Hom | # Hets | # nonRef | heterozygosity | % Het/genome | Ts/Tv | # Indels | ave depth | # Singletons |
| --- | --- | --- | --- | --- | --- | --- | --- | --- | --- | --- | --- | --- |
| 1A | 121,564,184 | 1563 | 2206 | 29,229 | 53,932 | 83,161 | 1.85 | 0.45 | 2.95 | 5,029 | 2091.3 | 128 |
| 1B | 121,021,253 | 1365 | 1718 | 28,528 | 55,200 | 83,728 | 1.93 | 0.46 | 2.95 | 5,085 | 2053.7 | 125 |
| 3 | 119,824,714 | 1192 | 1314 | 27,374 | 62,001 | 89,375 | 2.26 | 0.51 | 2.95 | 5,360 | 2110.3 | 14 |
| 4 | 114,826,852 | 1294 | 1508 | 28,211 | 60,152 | 88,363 | 2.13 | 0.50 | 2.96 | 5,344 | 2012.7 | 49 |
| 5 | 104,713,806 | 1280 | 1507 | 27,740 | 60,625 | 88,365 | 2.19 | 0.50 | 2.94 | 5,286 | 1819.5 | 36 |
| 6 | 114,420,326 | 1147 | 1205 | 27,921 | 61,022 | 88,943 | 2.19 | 0.51 | 2.95 | 5,336 | 2015.9 | 25 |
| 7 | 118,684,945 | 1414 | 1722 | 28,954 | 57,106 | 86,060 | 1.97 | 0.47 | 2.95 | 5,178 | 2020.1 | 50 |
| 8 | 6,809,404 | 4232 | 14786 | 29,671 | 45,786 | 75,457 | 1.54 | 0.38 | 2.96 | 4,384 | 115.3 | 6 |
| 10 | 116,034,541 | 13426 | 43980 | 25,956 | 59,139 | 85,095 | 2.28 | 0.49 | 2.97 | 5,091 | 2005.9 | 382 |
| 12 | 106,738,203 | 1669 | 2474 | 31,854 | 46,533 | 78,387 | 1.46 | 0.39 | 2.95 | 4,754 | 1853.8 | 87 |
| Granvin-1 | 46,222,242 | 994 | 1023 | 26,647 | 63,099 | 89,746 | 2.37 | 0.52 | 2.96 | 5,404 | 794.7 | 23 |
| Honindal-1 | 59,830,373 | 974 | 992 | 26,015 | 64,397 | 90,412 | 2.48 | 0.53 | 2.94 | 5,431 | 1026.6 | 51 |
| Honindal-2 | 47,534,306 | 1115 | 1163 | 27,750 | 53,168 | 80,918 | 1.92 | 0.44 | 2.96 | 4,923 | 785.4 | 131 |
| Laerdal-2 | 25,739,918 | 1626 | 2281 | 29,256 | 55,853 | 85,109 | 1.91 | 0.46 | 2.96 | 5,014 | 390.1 | 0 |
| Stordal-Ebbegarden-1 | 32,614,923 | 1502 | 1937 | 30,570 | 47,916 | 78,486 | 1.57 | 0.40 | 2.94 | 4,746 | 549.5 | 29 |
| Voss-1 | 57,745,958 | 954 | 966 | 27,337 | 62,322 | 89,659 | 2.28 | 0.52 | 2.95 | 5,383 | 1018.7 | 31 |
| Sums & Averages | 1,314,325,948 | 2234 | 5049 | 453,013 | 908,251 | 1,361,264 | 2.00 | 0.47 | 2.95 | 81,748 |  | 1167 |

**Table 3:** Top 10 genes by the number of high-impact variants. Variant impact as predicted by SnpEff

| Gene name | Gene Id | Description (from SGD) | # variants | # variants with impact |  |  |
| --- | --- | --- | --- | --- | --- | --- |
|  |  |  | missense | HIGH▽ | LOW | MODERATE |
| **FLO5** | YHR211W | Lectin-like cell wall protein (flocculin) involved in flocculation; binds mannose chains on the surface of other cells, confers floc-forming ability that is chymotrypsin resistant but heat labile; important for co-flocculation with other yeasts, mediating interaction with specific species; FLO5 has a paralog, FLO1, that arose from a segmental duplication | 44 | 18 | 56 | 47 |
| **MAL31** | YBR298C | Maltose permease; high-affinity maltose transporter (alpha-glucoside transporter); encoded in the MAL3 complex locus; member of the 12 transmembrane domain superfamily of sugar transporters; functional in genomic reference strain S288C | 35 | 10 | 32 | 39 |
| **PRM9** | YAR031W | Pheromone-regulated protein; contains 3 predicted transmembrane segments and an FF sequence, a motif involved in COPII binding; member of DUP240 gene family; PRM9 has a paralog, PRM8, that arose from a segmental duplication | 20 | 10 | 11 | 20 |
| **COS7** | YDL248W | Endosomal protein involved in turnover of plasma membrane proteins; member of the DUP380 subfamily of conserved, often subtelomeric COS genes; required for the multivesicular vesicle body sorting pathway that internalizes plasma membrane proteins for degradation; Cos proteins provide ubiquitin in trans for nonubiquitinated cargo proteins; the authentic, non-tagged protein is detected in highly purified mitochondria in high-throughput studies | 30 | 9 | 8 | 30 |
| **HXT4** | YHR092C | High-affinity glucose transporter; member of the major facilitator superfamily, expression is induced by low levels of glucose and repressed by high levels of glucose; HXT4 has a paralog, HXT7, that arose from the whole genome duplication | 5 | 8 | 22 | 5 |
| **YHL008C** | YHL008C | Putative protein of unknown function; may be involved in the uptake of chloride ions; does not appear to be involved in monocarboxylic acid transport; green fluorescent protein (GFP)-fusion protein localizes to the vacuole | 51 | 6 | 92 | 53 |
| **EHD3** | YDR036C | 3-hydroxyisobutyryl-CoA hydrolase; member of a family of enoyl-CoA hydratase/isomerases; non-tagged protein is detected in highly purified mitochondria in high-throughput studies; phosphorylated; mutation affects fluid-phase endocytosis | 50 | 6 | 106 | 50 |
| **YIR044C** | YIR043C | Possible pseudogene in strain S288C; YIR044C and the adjacent ORF, YIR043C, together may encode a non-functional member of the conserved, often subtelomerically-encoded Cos protein family | 28 | 6 | 18 | 28 |
| **YER186C** | YER186C | Putative protein of unknown function | 22 | 6 | 11 | 22 |
| **HXT1** | YHR094C | Low-affinity glucose transporter of the major facilitator superfamily; expression is induced by Hxk2p in the presence of glucose and repressed by Rgt1p when glucose is limiting; HXT1 has a paralog, HXT6, what arose from the whole genome duplication | 11 | 6 | 34 | 11 |

### Whole genome sequencing and assembly

Illumina paired-end sequencing resulted in 2.09 billion fragments and a total 277 billion bases. Raw reads have been deposited in the European Nucleotide Archive (ENA) under study accession PRJEB63528. Mapping rates of trimmed reads to the *S. cerevisiae* reference genome were > 99% for all our samples and 98% of all reads were properly paired. We generated *de novo* assemblies from trimmed reads for all kveik strains. Scaffold N50 was between 1,314 and 43,980 bases for our data and between 966 and 2,281 bases for the data from Preiss et al. (2018).

### Variant analysis

To better understand genotypic characteristics of kveik, we first investigated short nucleotide variants within all kveik strains resulting in 160,142 SNPs and 15,463 insertions or deletions (InDels) before any filtering (Supplementary Data S1) with a total of 6,552 multiallelic sites. 129,090 genotyped polymorphic sites remained after filtering (Supplementary Data S2), out of which 121,763 were SNPs (*ts/tv* = 2.95) and 7,327 InDels. Heterozygosity ratio (*nHet/nHom*) over all samples was 2.00 and varied only marginally between our and samples from Preiss et al. (2018) (1.97 and 2.07, respectively). Per-sample statistics of short variants detected are given in Table 2.

Genome-wide nucleotide diversity for all kveik strains was determined by ρε = 0.0039 (n=16) which is higher than for most *S. cerevisiae* subpopulations studied by Peter et al. (2018) except for the Mosaic region 3 population in their study (ρε = 0.0041924, n = 113); Tajima’s D = 1.716865 indicating scarcity of rare variants and increased heterozygosity.

We then predicted variant effects with respect to the reference assembly using SnpEff. Out of 6446 genes in the reference annotation, 671 (10%) genes had at least one variant with impact categorized as high by SnpEff and out of 6016 protein-coding genes, 5060 genes (84%) had at least one missense variant (Supplementary Data S3). Among the genes most affected by high-impact variants were the flocculation-associated gene FLO5 (YAR050W), two maltose associated genes from a subtelomeric complex locus MAL31 (YBR298C), and MAL33 (YBR297W), as well as genes encoding pheromone-regulated proteins PRM8 (YGL053W) and PRM9 (YAR031W).

Gene-ontology (GO) enrichment analysis of the 66 genes having at least three high-impact variants yielded 15 significantly enriched (p < 0.01) GO-terms from the biological process (BP) ontology (Supplementary Data S4) and 22 from the molecular function (MF) ontology (Supplementary Data S5). Most of the enriched terms are related to membrane transport of different sugars.

### Association analysis

To assess which variants are significantly associated with kveik and beer phenotypes, we performed association testing in of missense variants in PLINK. 66,378 loci were included in the analysis, out of which 18,889 loci were significantly associated with a group at p < 0.05 and 391 loci located in 185 different genes were significant after Bonferroni correction (Supplementary Data S6, S7). The five top-scoring loci, of which three localize to FLO9, were all perfectly segregating with homozygous non-reference alleles in all kveik strains and reference allele in all beer strains. Among the genes affected by the largest number of significant loci are MRC1, NUT1, FLO9 and MAL33 (Supplementary Dat S6). GO enrichment analysis of the 45 genes with at least three variants significant after correction did not yield significant terms from either MF or BP ontology.

### Variant effect analysis of selected heat shock-related genes

We wanted to investigate potential causative variants for the stress resistant phenotype of kveik in greater detail. We searched for genes annotated as involved in heat shock response that were affected by high impact variants and missense variants.

We manually investigated genotypes of selected genes with the largest number of missense mutations. Among the 26 heat shock-related genes with at least one amino-acid change, most had no high-impact (including disruptive) mutations, indicating high essentiality. Four genes had a single amino acid change and RPI1 had three such mutations. HSF1 (YGL073W) encoding Trimeric heat shock transcription factor, had the largest number of missense mutations (22). While 11 loci were significant before Bonferroni correction, none of the alleles showed a convincing genotype pattern upon inspection.

WSC3 (YOL105C) is a gene located on chromosome XV. We identified 16 missense variants and no disruptive variant in its coding region. 13 sites were significant in the association test, albeit not after Bonferroni correction. Of note, kveik strains contain several loci within WSC3 that have heterozygous non-reference alleles not evident in the investigated beer strains (Figure 4).

**Figure 4:**
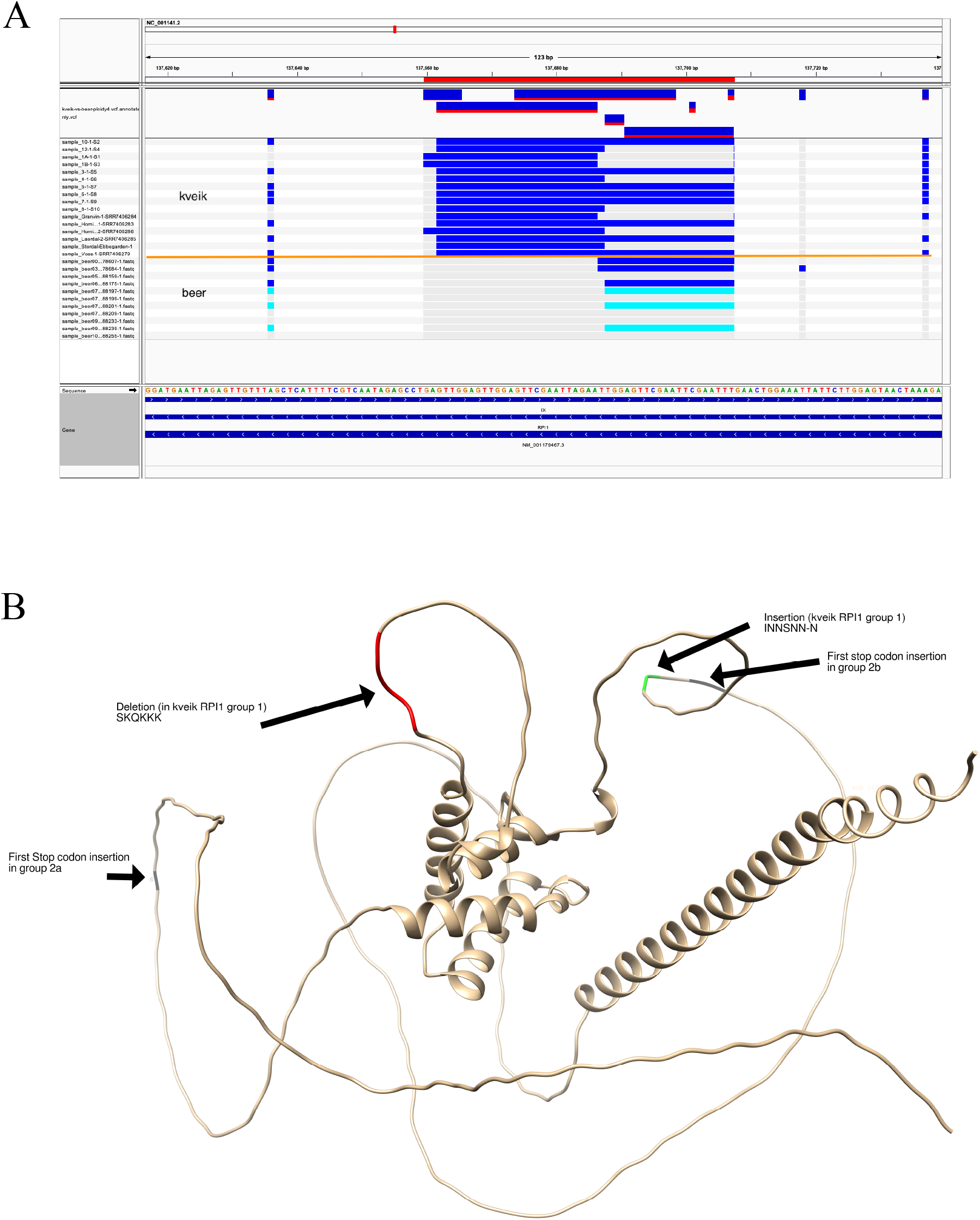
Visualization of variants in RPI1and predicted effects on the protein sequence. A) kveik-specific InDels over the genomic sequence of the RPI1 gene. Heterozygous variants are depicted by blue bands, homozygous variants by cyan bands. Kveik and beer groups are marked. B) 3D-model (predicted by AlphaFold) for RPI1 depicting localization of possible variants (stop-codons, InDels) derived from consensus genotypes.

**Figure 5:**
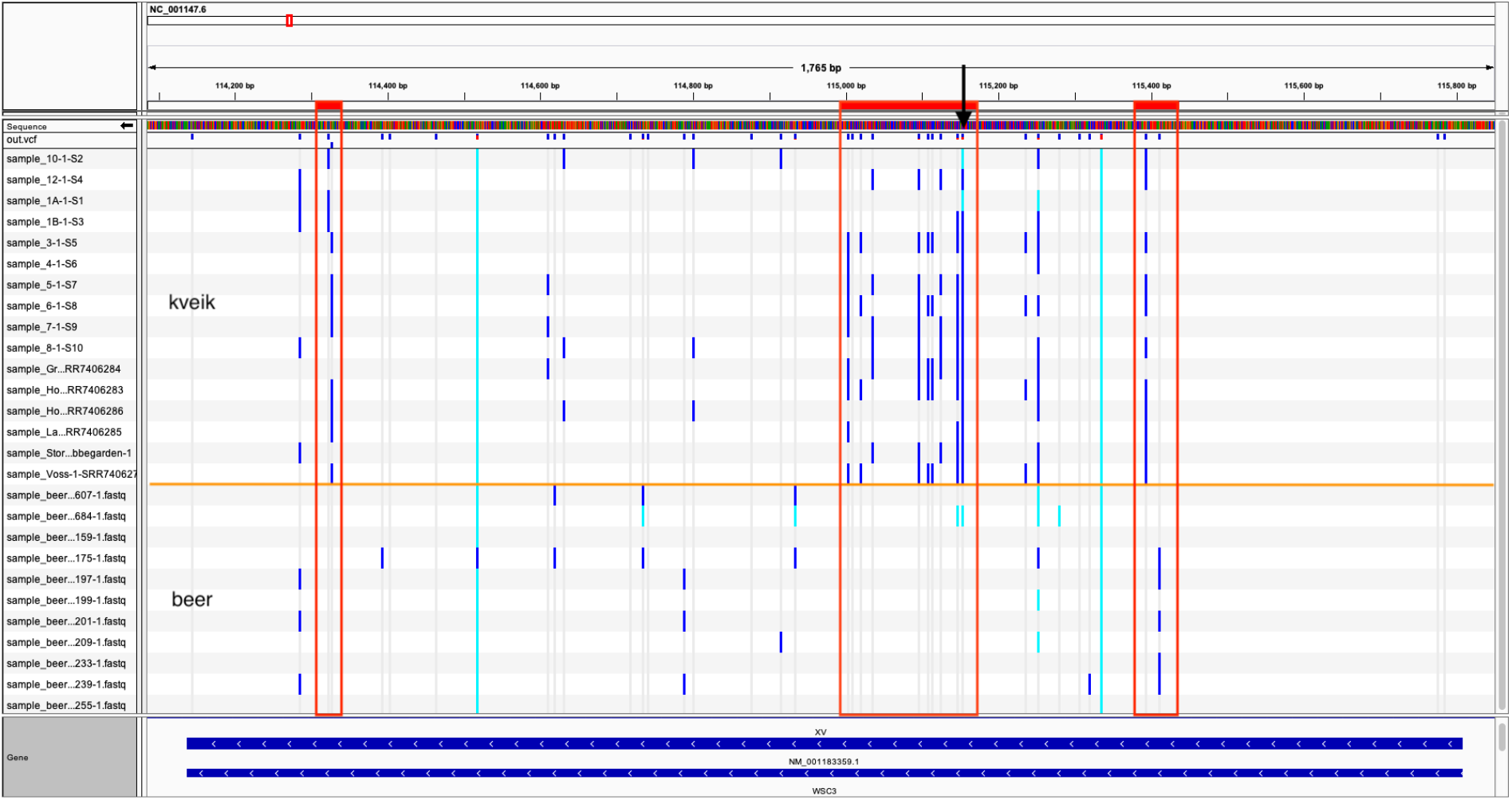
Kveik-specific missense variants in the coding region of the WCS3 gene. Vertical bands indicate polymorphic sites, grey: homozygous reference allele, cyan: homozygous ALT allele, blue: heterozygous allele. Regions of interest boxed. Arrow: location of kveik-specific missense allele c.655T>A|p.Ser219Thr which is significantly associated with the groups (p = 1.01e-05). Visualization shows unfiltered tetraploid variants.

Overall, only five heat shock-related genes had at least one variant classified as high impact within the filtered variants (RPI1, MGA1, GAC1, SSE2, ECM10). We focus here on RPI1 (YIL119C). RPI1 is a gene located on chromosome IX encoding a transcription factor. The gene sequence RPI1 contains a region enriched for larger heterozygous InDels which were found in kveik but not in beer strains (Figure 3A).

To assess the effect of eventual stop codon insertions on the protein sequence, we generated consensus sequences of the RPI1 gene from the aligned reads for each kveik strain and translated them. In a multiple sequence alignment, conservation of the amino-acid sequence declined massively after residue 60 (Supplementary Figure 1). The obtained genotypes can be roughly divided into two groups: 1) 7 samples with intact amino acids sequence without premature stop codons (from samples 1A, 1B, 4, 5, 8, Hornindal 2, and Stordal Ebbegården 1); 2) 9 samples with premature stop codons (from samples 3, 6, 7, 10, 12, Granvin 1, Hornindal 1, Laerdal 1, and Voss 1). Group 2 can be further subdivided by approximate residue of the first stop codon (group 2a and 2b in Supplementary Figure 1). Two pairs of samples obtained from the same provider (1A & 1B, Laerdal 1 & 7) fall into the same major groups. For the intact sequences in group 1, all members are highly identical ([95.7%, 100%]), apart from minor short indels in three low-complexity regions at residues 60, 170, and 251. Figure 3B depicts the approximate sites of mutations on the predicted 3D-structure of the gene product. All variants are in loop regions or disordered regions of the protein. At least for stop-codon insertions closest to the N-terminus (from residue 60), the protein function is likely disrupted.

### Phylogenetic analysis based on de novo assemblies

To generate a Maximum Likelihood (ML) phylogenetic tree, we aligned a total of 179 *S. cerevisiae* genomes including our assemblies (represented by red dots in Figure 1), six kveik strains previously reported (Preiss et al. 2018) (represented by blue dots in Figure 1), 157 domesticated and wild strains reported by (Gallone et al. 2016), five wild *S. cerevisiae* isolates from primeval Chinese forests (Duan et al. 2018), and the *S. paradoxus* reference genome to root the tree.

Phylogenetic analysis showed that the kveik isolates from both studies formed a compact monophyletic group with no signs of batch effects, as seen in both PCA and the ML tree (Figure 2A, 3A, Supplementary Figure 2). Choice of assembly method (IDBA-UD assembler in the reference genomes by Gallone et al. (2016) vs. SPAdes in our de-novo assemblies) had marginal to no impact on placement and branch-length. Within the phylogenetic tree, kveik presented as a pure sister clade with 100% bootstrap support to a large group consisting predominantly of domesticated yeasts. Within the Kveik clade itself, strains clustered predominantly by geographic origin, with a notable cluster from the Voss region (Figure 1B, 2A). The largest deviation from clustering by geographic origin was found for sample Hornindal-1 which – unlike sample Hornindal-2 - did not group with the samples from the same area but was phylogenetically closer to samples from the more southern regions of Laerdal and Granvin. Of note, samples from the same owner at different time-points (samples 1A and 1B; sample 6 and Voss-1; and sample 7 and Laerdal-2) clustered closely together.

When rooted on the *S. paradoxus* branch, the wild samples from primordial Chinese forests appeared closest to the root of the *S. cerevisiae* tree as previously reported. Meanwhile, the other wild samples from the study of Gallone et al. (2016) formed a sister group to sake and bioethanol strains from the same study, indicating an early domestication of these strains.

### Phylogenetic analysis based on variants

We performed a series of computational experiments generating phylogenetic trees based on variant calls to confirm the phylogenetic position of kveik obtained from the de novo assemblies. There, we treated all samples in an identical way and used variant calls by the GATK pipeline to generate consensus genotypes for a subset of 41 selected kveik, wild, and domesticated *S. cerevisiae* strains. We then generated phylogenetic trees for the consensus genotypes consisting either solely of the reference alleles (REF) or the alternative alleles (ALT) and filtered at different MAF thresholds (0.01, 0.05, 0.1, 0.2, and 0.3) and in one experiment retained only variants annotated overlapping a protein coding gene region.

From 41 samples and before filtering, 283,791 SNPs and 25,961 indels were recovered at a total of 307,789 polymorphic sites. Kveik and wild strains also appeared to be more similar to the reference sequence than domesticated strains. Tree topology was highly variable and depended mostly on whether REF or ALT alleles were used for the consensus genotype and secondarily also on the MAF threshold used for filtering. Domesticated strains had much larger branch lengths than kveik strains as expected from the larger number of variants. In none of the primary phylogenies, nor generated consensus tree (Figure 3B, Supplementary Figure 3) did the kveik strains form a subgroup within the domesticated groups. Notably, trees did never place a common ancestor of kveik within the samples from the Beer I group in our analysis. In most cases, kveik samples presented as a sister group to beer yeasts.

### Assessment of phylogenetic analysis using a mixture of different consensus calling methods

To compare our phylogenetic analyses with previous studies, we conducted calibration analyses integrating different consensus calling methods. We generated consensus sequences for all kveik samples based on filtered variant calls, while introducing all other sequences as genome assemblies. We also downloaded the raw sequencing data of randomly selected samples from Gallone et al. (2016) and subjected them to our implementation of the variant calling pipeline used by Preiss et al. (2018). The derived consensus calls were then added to the multiple sequence alignment as calibration data and a ML phylogeny was computed. We expected that if there was no method bias, all calibration sequences should cluster with their respective genome assembly in the phylogenetic tree.

The resulting tree topology was similar to the original study, with the kveik samples forming a compact group within other mainstream brewing strains, albeit with inflated branch lengths. We repeated the analysis with two additional datasets and each time the consensus sequences integrated in a different location in the tree (Supplementary Figure 4A-C) but never clustered within the Beer I group or close to taxa beer072, beer074, beer093. We observed that calibration samples tended to cluster together instead of with their respective genome assembly in almost half of the cases for the largest dataset, as depicted by blue branches and red asterisk in Supplementary Figure 4B and 4C. This indicates significant bias attributed to the consensus methods used and lack of robustness.

### Yeast strains in kveik samples after reuse in brewing

Previous work indicates that individual kveik yeast cultures show diverse colony morphology which may reflect strain diversity (Preiss et al. 2018). Thus, we isolated single colonies from kveik 1 two times after two years and reuse in brewing (Table 1). The two isolates (1A and 1B) show almost identical sample statistics of variant calling and de novo assembly (Table 2), and the two samples cluster closely together in phylogenetic analyses. In addition, two kveik types from our work are collected from the same location and owner as in the previous work by Preiss et al. (2018). Furthermore, these single colonies picked at different times in different laboratories show minor genetic differences and cluster closely together after several years of active use in brewing.

## Discussion

Our results call the previously determined phylogenetic relationship of kveik as a compact sub-clade of other domesticated *S. cerevisiae* strains into doubt. While Preiss et al. (2018) hypothesized a common ancestor of the kveik strains within a paraphyletic Beer 1 clade of conventional brewing yeast, in our analysis, kveik presents as a monophyletic sister group to most other domesticated *S. cerevisiae* strains. The latter tree topology thus places the origin of kveik closer to the root of the *S. cerevisiae* tree than previously thought and we argue that such a topology is more credible for reasons we discuss in the following.

By performing a series of calibration experiments we noted that a mixed approach for generating a variant matrix from two different consensus calling methods, namely *de novo* assemblies and *de novo* variant calling, failed. Evidently, there is a tendency of otherwise unrelated taxa to cluster based on the method of variant calling. We interpret these clusters as artifacts likely caused by combining incomparable methods.

To better understand the reasons behind this assumption, it is relevant to take a closer look at the methods employed to generate consensus sequences and how they are fundamentally different. The reference genomes from (Gallone et al. 2016) were all assembled using the IDBA-UD algorithm (Peng et al. 2012). DeBruijn Graph (DBG) assemblers, such as IDBA-UD and SPAdes (Bankevich et al. 2012), employ error correction steps at various stages of the assembly, including removal of “bubbles” in the graph that may be introduced by sequencing errors or sequence variants. Implementation details differ slightly between software, however, when applying bubble removal to the DBG, assemblers generally give preference by coverage. It is therefore reasonable to assume that a polymorphic site will likely be represented by the allele with highest coverage in the consensus sequences produced by a DBG assembler. On the other hand, the variant calling pipeline based on Freebayes and BCFtools has a very different focus. In particular, the consensus method provided by BCFtools, will by default (and as applied by Preiss et al. (2018)) report the ALT (non-reference) allele, irrespective of coverage, frequency or other statistics.

This train of thought also explains inflated evolutionary rates of kveik observed in the paraphyletic Beer 1-kveik clade when using a mixed consensus analysis (Fig.2 in Preiss et al. (2018)). Branch length heterogeneity may distort the phylogeny and lead to random clustering of otherwise unrelated taxa, also known as long-branch attraction (Felsenstein 1978). We propose further that these differences in evolutionary rates, which violate the stationarity assumption, are not real but were introduced by using different consensus methods. Therefore, the previous placement of kveik within the Beer 1 group by Preiss et al. (2018) is likely an artifact. We conclude that only consensus generating methods of the same type, either variant analysis or *de novo* assemblies should be combined in one tree to avoid such distortions by incompatible rates. To further support this claim, we also constructed phylogenetic trees based on variant analysis only. Important conclusions arise from these experiments: Firstly, in our analysis we never saw kveik placed even close to the clade it had been placed in previously, irrespective of the method used. Secondly, tree topologies based on *de novo* variant calls are heavily influenced by allele selection criteria and therefore appear less robust than *de novo* assembly.

The genetic diversity of the tetraploid and highly heterozygous kveik yeasts was only slightly higher than among other *S. cerevisiae* isolates (*π* = 3 × 10^−3^) (Peter et al. 2018), but considerably higher than the diploid sake-yeasts that show low heterozygosity (Duan et al. 2018). Numbers of polymorphic sites reported tend to vary slightly between studies, e.g.: 121,763 SNPs (this study, after hard filtration) versus 142,120 loci reported by (Preiss et al. 2018) using a different method including more strains. However, the accuracy of variant calls may depend on multiple factors such as the number of samples included, software, read depth, and filtration parameters (Barbitoff et al. 2022).

For three of the kveik-types genetic changes after 2 years or more with regular reuse in traditional farmhouse brewing were minor. Genomic analyses of multiple kveik strains of the same origin have not been performed elsewhere. Preiss et al. (2018) reported differences in colony morphology and DNA fingerprinting at 30°C for different strains from the same kveik yeast culture. We propose that kveik yeast may occupy a niche under high selective pressure from traditional brewing practices including high brewing temperature where the yeasts are harvested and re-used after the fermentation process as described for beer yeasts (Gallone et al. 2016).

High temperature- and ethanol tolerance has been observed previously in a large proportion of kveik yeast strains and such phenotypes may be favorable for brewing as well as bioprocessing applications (Preiss et al. 2018). For yeast and other organisms, it has been long known that responses to different stress conditions, and heat and ethanol resistance in particular (Piper 1995), are linked and can be governed by cross-resistance at different cellular levels by inducing either specific or general stress response mechanisms (Swiecilo 2016). However, little is known about the causative genes and variants enabling the specific stress tolerance found in kveik. Classical heat-shock proteins, e.g., HSP104 are required for survivability under and respond to a multitude of stress factors (Sanchez et al. 1992). All genes of this family had none or few coding variants and no high impact variants whatsoever in kveik strains.

A focus has previously been on observed loss-of-function mutations such as frame-shift mutations. Genes such as KEX1, LRG1, SWP82, RPI1, IRA1/IRA2, and CDC25 have been named in previous work. In our analysis, we did not always detect the same loss-of-function variants e.g., KEX1, LRG1, SWP82, and IRA1/2 where we detected several significant missense variants but no disruptive variant at all (KEX1) or these variants did not pass our filtering criteria. However, and notwithstanding differences in analysis procedures and their accuracy, to propose a possible association of variants with the kveik phenotype more in-depth analyses of their distribution across strains and effects of respective null mutants are necessary.

RPI1 is a gene encoding a transcription factor that acts as a negative regulator on the RAS cAMP pathway (Kim and Powers 1991). It has been demonstrated that RPI1 is important for stress tolerance under ethanolic fermentation at high temperature (Puria et al. 2009). In accordance with Preiss et al. (2018) we found strong evidence for the presence of heterozygous frame-shift variants that are likely to disrupt protein function in some kveik strains following from our structural analysis. It is important to note, however, that in genetics studies the stress-tolerant phenotype was induced by overexpression of RPI1 and that the null mutant rpi1-Δ had lower resistance to heat and other stressors (Brown et al. 2006; Jarolim et al. 2013; Kim and Powers 1991; Mira et al. 2009; Puria et al. 2009). Phenotypes of null-mutants of other genes, namely lrg1-Δ and ira1/2-Δ showed a similar tendency of increased stress sensitivity (from SGD) or were inconclusive. We thus hypothesize that the frame-shift variants in RPI1 are not causative for heat tolerance directly. One may speculate that there exists a dose-dependent fitness advantage of lower RPI1 expression during the life cycle of kveik, but then haplotypes encoding a functional RPI1 would remain under strong selection against homozygous loss-of-function. To assess this further though, we would need a larger number of strains and properly phased haplotypes.

We further focus on missense variants. However, 84% of all genes are affected by at least one missense variant, thus requiring another filtering step. As kveik and beer form distinct clades in our phylogenetic analysis, the question arises which sites most specifically segregate these groups, and thereby might contribute to phenotypic differences. With now 16 isolates at our disposal, it was possible to do a relatively simple association test to determine variants significantly associated with the complex kveik phenotype but not individual traits. Among the genes harboring the highest number of highly significant variants were flocculation associated genes (FLO9, FLO5) and a protein involved in DNA replication and repair under stress (MRC1).

Transcriptional regulation of the heat shock response in *S. cerevisiae* is governed by the transcription factors HSF1, MSN2, and MSN4. These genes did not have any high-impact mutation, but many amino-acid changes were detected hinting at a regulatory rather than structural adaptation of the stress response pathways. Several of the mutations in HSF1 and MSN2/4 were also mildly significantly associated with the kveik-versus-beer phenotype. However, upon visual inspection none of the alleles was specific for either group.

Accumulation of the disaccharide trehalose has been known to contribute to cellular stress resistance (Jain and Roy 2009). Recently, Foster et al. (2022) reported a significant increase in intracellular trehalose accumulation and proposed mutations in the Trehalose Synthase Complex. The candidate variant 2500A>G (Ser834Gly) in the coding sequence of TPS3 was found to be homozygous in all kveik strains. We identified the same missense variant, and it was also the only variant in the TPS genes with genome-wide significant association. The variant was homozygous also in our strains, except for 1A and 1B where it was heterozygous.

Among variants that might contribute to stress-tolerance of kveik we propose a group of heterozygous missense variants in the WSC3 gene (Sensor-transducer of the stress-activated PKC1-MPK1 signaling pathway) as the best candidates based on our association analysis. The WSC genes SLG1/WSC1, WSC2, and WSC3 are integral membrane proteins that are involved in response to heat and other stressors and maintenance of cell-wall integrity (Verna et al. 1997). WSC null mutants were found to be more sensitive to heat-shock, ethanol and other stressors (Zu, Verna, and Ballester 2001). In accordance with this, none of the WSC genes contained a high-impact variant in any kveik strain.

It has been demonstrated that the phylogenetic position of the solid-state fermentation group from China and Far East Asia (e.g., the Japanese Sake and the Chinese Huangjiiu lineages) is close to the center of origin of domesticated *S. cerevisiae* (Duan et al. 2018). In addition, as for kveik some of these solid-state fermentation strains grow well at 41°C (Duan et al. 2018). Both from a phenotypic and genomic perspective it is therefore tempting to speculate on an evolutionary link between these Asian rice and barley fermenting yeasts and the Norwegian kveik beer yeast.

One may argue that inducible temperature and ethanol tolerance have been observed in a multitude laboratory and wild strains. Hence, high conservation of genes involved in stress tolerance rather than presence of any singular mutation may contribute to this phenotype in kveik. In the context of brewing, however, it is not only survivability or the mere ability to ferment that is relevant. At these high temperatures, many commercial strains would often produce an excess in off-flavors such as fusel alcohols (Kits and Garshol 2021). Therefore, a more complex and specific definition of the kveik phenotype comprises that it not only thrives but also maintains a metabolite composition resulting in fermentation products favorable to human consumption.

The biosynthesis of fusel alcohols is via the Ehrlich pathway named after Felix Ehrlich who first proposed that “fusel oils” were derived from aromatic amino acids (Ehrlich 1907). The Ehrlich pathway comprises a series of enzymatic reactions involving - among others – an array of alcohol- and aldehyde dehydrogenases and related transporters and regulatory genes (reviewed in Hazelwood et al. 2008). Because of the purported reduction in off-flavors in kveik beer, we deemed this pathway an interesting candidate for further investigations. In our data, missense variants were most common in regulatory genes rather than in genes encoding enzymes in this pathway. Namely the transcription factors ARO80 and ADR1 had most missense variants but alleles were not very specific to kveik (Supplementary Data S8). AAD4 is a putative aryl-alcohol dehydrogenase also involved in the oxidative stress response (Delneri, Gardner, and Oliver 1999). The coding gene contains several highly significant and predominantly homozygous missense variants and homozygous frame-shift variants.

Due to the lack of quantitative data on phenotype-genotypes associations in this group, further studies on strain variation associated with stress tolerance using a larger number of kveik and conventional beer strains, as well as solid-state fermenters are justified. Analysis of quantitative trait loci (QTL) could, for example, improve our knowledge on how kveik is adapted to its specific ecological niche set by the historical tradition of farmhouse brewing. Nevertheless, QTL analyses often require one order of magnitude more samples than we currently have available.

Overall, our results indicate that the kveik yeast samples form a compact monophyletic group close to the root of domesticated *S. cerevisiae*. This finding is comparable to previous studies on the origin of the Sake yeast group. Separate domestication events for sake and wine have been proposed (Fay and Benavides 2005; Liti et al. 2009) and China or Far east Asia may be the origin of domesticated *S. cerevisiae* (Bing et al. 2014; Duan et al. 2018; Peter et al. 2018; Wang et al. 2012). Thus, our results support an ancient origin also for kveik yeasts, which by necessity implies that the domestication of kveik took place far earlier than previously anticipated. Considering a possible far eastern or eastern European origin, the apparent endemism to western Norway remains as a big paradox, which is not resolved by this study. However, as a domesticated yeast lineage, kveik depends on traditional farmhouse brewing, which is vastly abandoned. Consequently, sampling for kveik or an elusive missing link, whether extant or extinct, needs to pinpoint these relevant environments and substrates where farmhouse brewing has taken place recently.

It is not known whether kveik is still present in specific environments in the historic long-distance trade network of western Norway, or elsewhere along the hypothetical migration route from Asia to Europe (e.g., along the Silk Road) in places where the farmhouse brewing culture has been maintained. To investigate this hypothesis further, metagenomes from historic or ancient objects used in fermentation of beer and other products should be obtained.

## Acknowledgements

We thank the kveik owners for sharing the yeast with us for this research, and Jorunn Børve, Lorena Butinar and Ingunn Øvsthus for advice and assistance. The work is financed in part by a grant from the Regional Research Council of western Norway and in part by research funding from Western Norway Culture Academy and NIBIO. The Genomics Core Facility (GCF) at the University of Bergen, which is a part of the NorSeq consortium, provided services on the project; GCF is supported in part by major grants from the Research Council of Norway (grant no. 245979/F50). Bioinformatics analysis was carried out under the framework of ELIXIR Norway.

## Data availability

Raw sequencing reads have been submitted to ENA under study accession PRJEB63528. Supplementary data and code have been deposited in figshare under <u>DOI: 10.6084/m9.figshare.c.6724260.v1</u>.

## Author contributions

HGE, AOE and MD: study design. HGE, AOE, AE, DD and MD: sample and data collection. HGE, LKH, AE and RH: laboratory sample analysis. MD, DD, HGE and SG: bioinformatic analysis and data analysis. HGE, MD, AOE, LKH, DD, SG, SH, AE and TM: manuscript writing and editing. All authors contributed to the article and approved the submitted version.

## Notes

https://doi.org/10.6084/m9.figshare.c.6724260.v1

### Competing Interest Statement

The authors have declared no competing interest.

